# A severely affected MRL/Lpr mouse exhibits a divergent clinical–immune profile associated with a profound metabolic alteration

**DOI:** 10.64898/2026.01.07.698139

**Authors:** Karim Matmat, Carole Jamey, Daniel Brumaru, Ayikoé-Guy Mensah-Nyagan, Hélène Jeltsch-David

**Affiliations:** Inserm – Centre d’Investigation Clinique 1434, Neuroprotection and Remyelination – Centre de Recherche en Biomédecine de Strasbourg, University of Strasbourg, Strasbourg, France; Laboratoire de Biochimie et de Biologie Moléculaire, Pôle de biologie, Hôpitaux Universitaires de Strasbourg, Strasbourg, France; Laboratoire des Sciences de l’Ingénieur de l’Informatique et de l’Imagerie ICube, UMR 7357, CNRS / Université de Strasbourg / INSA / ENGEES / INRIA, Strasbourg, France

## Abstract

Systemic lupus erythematosus (SLE) displays marked clinical and biological heterogeneity that is incompletely captured by group-based analyses. Although the MRL/Lpr mouse is a well-established lupus model, individual-level divergence within this genetically homogeneous strain remains poorly characterized.

Here, we describe an MRL/Lpr mouse exhibiting an exceptionally severe systemic phenotype and provide an integrative characterization combining clinical assessment, inflammatory profiling, neuroaxonal injury, and targeted metabolomics. Despite pronounced clinical deterioration, including severe proteinuria, reduced organ weights, and marked neuroaxonal damage, this mouse did not show a globally exacerbated cytokine profile relative to other MRL/Lpr animals. In contrast, plasma neurofilament light chain levels were massively elevated, indicating substantial neuroaxonal injury. Targeted metabolomic analysis revealed a profoundly altered biochemical signature, with coordinated disruptions in nitrogen handling, sulfur amino acid metabolism, and neurometabolic pathways, clearly separating this animal from both control and lupus-prone peers.

These findings illustrate that extreme disease severity can emerge independently of overt cytokine escalation and identify metabolic dysregulation as a major dimension of pathological divergence at advanced disease stages. Although descriptive and based on a single individual, this work highlights the value of extreme phenotypes for uncovering biological inflection points that remain concealed in averaged analyses and supports the integration of metabolic readouts alongside inflammatory markers in autoimmune disease research.

## Introduction

Systemic lupus erythematosus (SLE) is a heterogeneous autoimmune disorder marked by fluctuating inflammatory activity, multisystem involvement, and substantial inter-individual variability in disease trajectories [1–3]. The MRL/MpJ-Faslpr (MRL/Lpr) mouse strain is one of the most established spontaneous models of systemic autoimmunity, recapitulating key features such as lymphoproliferation, glomerulonephritis, neurobehavioral alterations, dermatitis, and chronic inflammation [4, 5]. Despite its genetic homogeneity this model nonetheless displays notable variability in disease onset and severity, thereby mirroring essential aspects of human SLE heterogeneity [6, 7]. Such intra-strain divergence is rarely explored at the individual level, although unusually severe presentations can reveal mechanisms undetectable in group-averaged analyses [8]. This is particularly relevant in chronic autoimmune disease, in which inflammatory burden, metabolic stress, and tissue injury may diverge substantially between individuals [9].

In this Short Communication, we describe an MRL/Lpr mouse exhibiting an exceptionally severe systemic phenotype. By integrating clinical, inflammatory, and metabolic assessments, we illustrate how disease severity can emerge independently of cytokine escalation.

## Materials & Methods

### Animals and pathological assessment

Female MRL/Lpr and congenic MRL+/+ mice (The Jackson Laboratory, US) were housed under standard conditions with *ad libitum* access to food and water. At 17-weeks of age, corresponding to the peak of systemic autoimmunity, animals were anesthetized with isoflurane and euthanized by decapitation. Blood was collected immediately and plasma was obtained by centrifugation (2,000 g, 5 min). Kidneys, spleen, brain, and spinal cord were rapidly dissected and weighed. Terminal proteinuria was assessed using Albustix® reagent strips (Siemens). One MRL/Lpr mouse (*#15*) displayed an unusually severe systemic phenotype and was therefore examined descriptively as an extreme case. All procedures complied with EU Directive 2010/63/EU and were approved by the institutional ethics committee (APAFIS#35144).

#### Cytokine and NfL quantification

Plasma cytokines levels were quantified using a custom U-PLEX multiplex assay (Meso Scale Discovery, MSD®) and neurofilament light chain (NfL) was measured using a U-PLEX singleplex kit from the same manufacturer. EDTA plasma samples were processed according to the manufacturer’s instructions, and electrochemiluminescence signals were acquired on a MESO QuickPlex SQ 120. Concentrations were interpolated from standard curves and expressed in pg/mL.

#### Targeted metabolomic profiling

Plasma amino acids and related metabolites were quantified using the MassChrom® Amino Acids kit (Chromsystems), a standardized clinical LC-MS/MS method, following the manufacturer’s instructions. Final concentrations (µmol/L) were used for downstream analyses.

#### Statistical analysis

Normality was assessed using the Shapiro–Wilk test and variance homogeneity with Levene’s test. Depending on these assumptions, comparisons were performed between MRL+/+ and MRL/Lpr mice using an unpaired t-test, Welch’s t-test, or a Mann–Whitney test. PCA was computed on scaled cytokine or metabolite data. The composite inflammatory score was generated by z-normalizing each cytokine. Mouse #15 was included in the MRL/Lpr group for all statistical analyses; its profile was then described qualitatively to illustrate intra-group variability. Analyses were performed in R (v4.3).

## Results

### Clinical phenotype reveals a severely affected MRL/Lpr mouse

At terminal assessment, MRL/Lpr mice showed the expected systemic alterations relative to MRL+/+ controls. Body weight was broadly comparable within the MRL/Lpr group, although mouse *#15* exhibited the lowest weight of all animals (Figure 1a). Kidney, spleen, and spinal cord weights were significantly increased in MRL/Lpr mice, along with elevated proteinuria (Figure 1b–c). Interestingly, mouse *#15* displayed an atypical organ pattern: kidney weight remained within the typical MRL/Lpr range, but spleen, brain, and spinal cord weights were the lowest of the group, together with a maximal proteinuria score (5/5). This animal also showed severe ulcerative dermatitis with complete loss of snout fur, associated with a visibly deteriorated general condition (Figure 1d).

**Figure 1.**
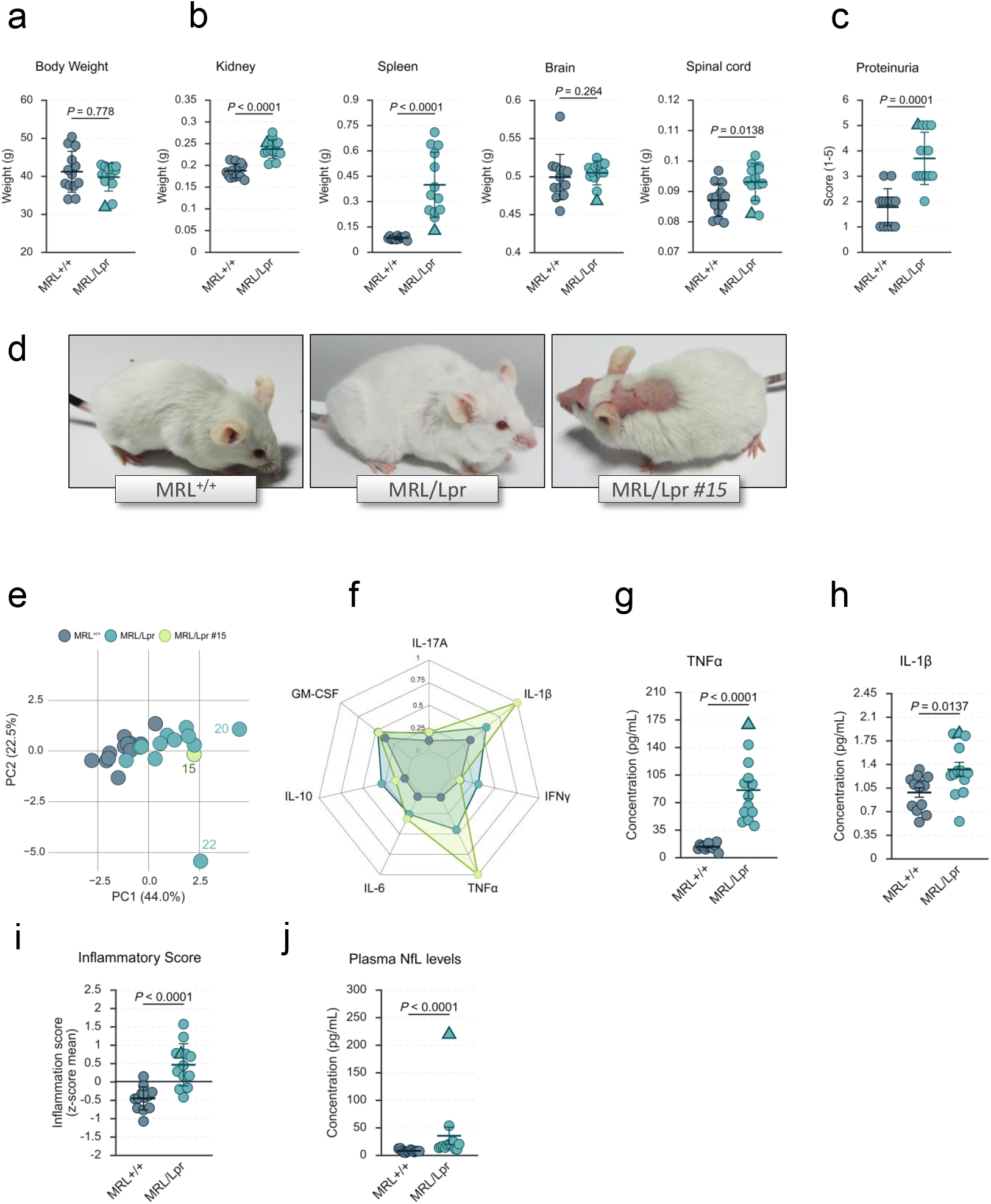
Systemic and behavioral profile of MRL/lpr mice at 17 weeks, illustrating the markedly aggravated condition of mouse *#15*. (a–c) Body and organs weights, and terminal proteinuria in MRL^+/+^ and MRL/lpr mice. Brain and spinal cord weights are included to document central nervous system tissue mass at endpoint. **(d)** Representative photographs showing the external phenotype of an MRL+/+ mouse, a typical MRL/lpr mouse, and mouse *#15* at sacrifice. **(e)** Principal component analysis (PCA) integrating all measured inflammatory cytokines. **(f)** Radar plot summarizing cytokine levels across groups, with the profile of mouse #15 highlighted. **(g-h)** Circulating concentrations of TNFα (g) and IL-1β (h). **(i)** Composite inflammatory score derived from normalized cytokine values. **(j)** Plasma neurofilament light chain (NfL) levels as a marker of neuroaxonal injury. Data are presented as individual values with mean ± SEM. Statistical procedures are detailed in the Methods section. Across all scatterplots, mouse *#15* is represented as a triangle and all other individuals as circles. Data are presented as individual values with mean ± SEM. Statistical procedures are detailed in the Methods section.

#### Inflammatory profiling shows limited cytokine dysregulation in mouse #15, but pronounced neuro-axonal injury

Principal component analysis (PCA) of plasma cytokines did not separate mouse *#15* from the remainder of the MRL/Lpr group (Figure 1e), indicating that its marked clinical deterioration was not associated with an overtly exaggerated cytokine profile. Other animals, particularly *#20* and *#22*, exhibited more divergent inflammatory signatures. Nevertheless, the radar plot revealed selective increases in TNFα and IL-1β in mouse *#15*, with values exceeding the MRL/Lpr group mean and ranking among the highest in the cohort, notably, TNFα reached the maximum recorded level (Figure 1f–h). Consistently, the composite inflammatory score confirmed this pattern, placing mouse *#15* in the upper range of group variability without exceeding the expected dispersion (Figure 1i).

In contrast, plasma NfL levels showed a striking divergence: mouse *#15* displayed a massive elevation, far surpassing all other animals (Figure 1j). This divergence indicates that neuro-axonal injury can escalate independently of systemic cytokine levels.

### Metabolic profiling reveals a profoundly altered signature in mouse #15

Targeted metabolomics showed a strongly altered biochemical profile in mouse *#15*. PCA clearly isolated this animal from both MRL+/+ controls and the remaining MRL/Lpr mice, indicating a global shift in amino-acid homeostasis (Figure 2a). This separation reflects a coordinated shift across multiple metabolites rather than isolated abnormalities. Fold-change analysis (*e.g*., |log_2_FC| > 2) identified major elevations in citrulline, alanine, and 3-methylhistidine, reflecting disruptions in nitrogen handling and muscle-derived metabolites, together with marked increases in cystathionine and taurine, indicative of altered sulfur amino-acid metabolism (Figure 2b–c). Conversely, several metabolites were markedly decreased, including a near-complete depletion of GABA. Pathway enrichment of metabolites meeting the |log_2_FC| > 2 threshold highlighted significant over-representation of the urea cycle, glutamate metabolism, ammonia recycling, and related nitrogen-processing pathways (Figure 2d).

**Figure 2.**
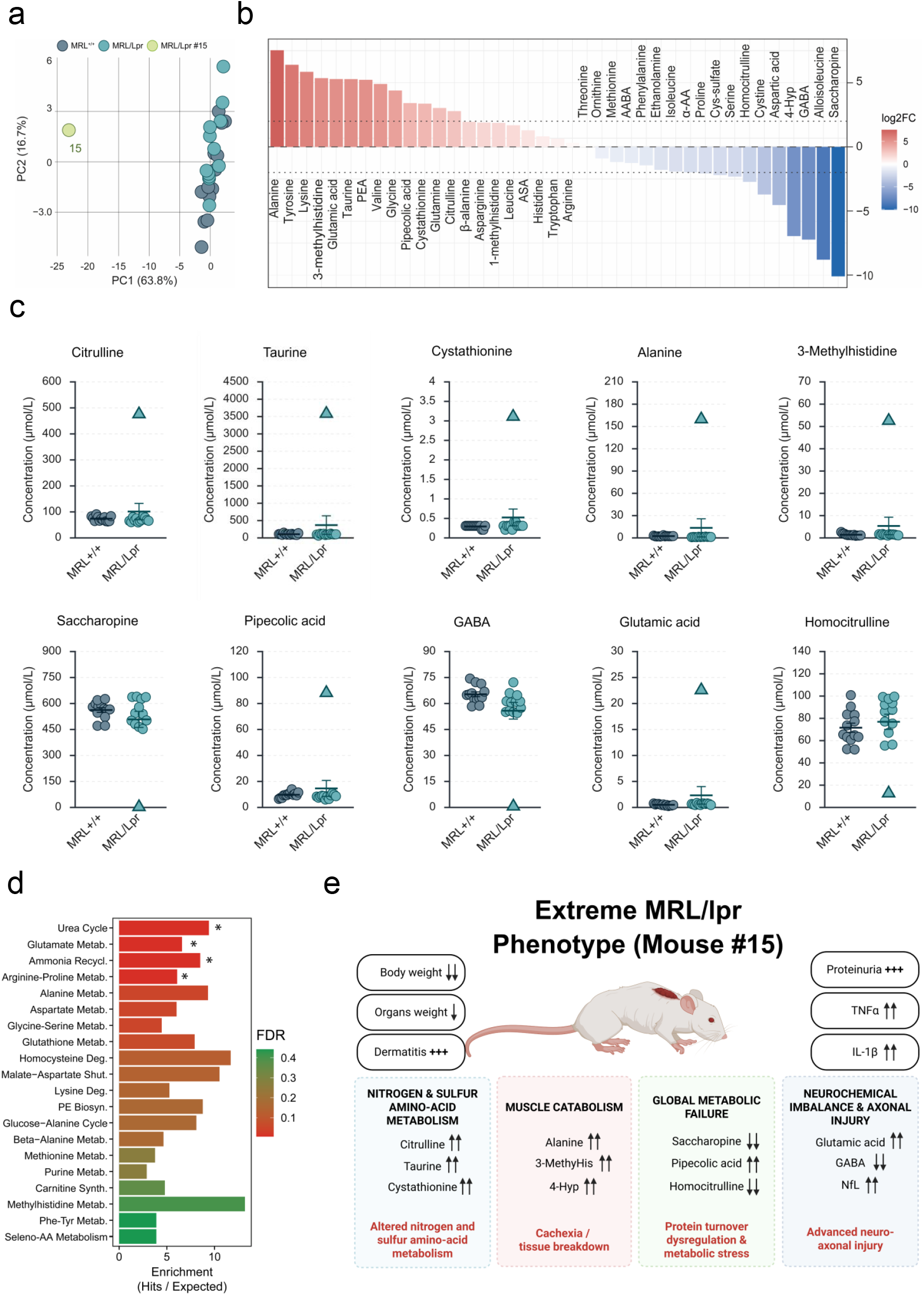
Metabolic profiling reveals a distinct biochemical signature in mouse #15. **(a)** PCA of plasma metabolites in MRL+/+ and MRL/lpr mice, highlighting the isolated metabolic position of mouse *#15*. **(b)** Log_2_ fold-change distribution of all quantified metabolites comparing mouse *#15* with the remaining MRL/lpr mice, illustrating the amplitude and direction of metabolic deviations. **(c)** Representative individual metabolite concentrations, including markers of nitrogen handling (urea cycle), sulfur amino-acid metabolism, muscle catabolism, and neurochemical imbalance. Mouse *#15* is displayed as a triangle across all scatterplots. **(d)** Pathway enrichment analysis based on altered metabolites, highlighting over-represented metabolic pathways (FDR-adjusted). **(e)** Integrative schematic summarizing the systemic phenotype of mouse #15, linking major clinical features with the corresponding metabolic alterations. Data are shown as individual values with mean ± SEM. Statistical procedures are described in the Methods.

Together, these alterations delineate a profoundly dysregulated metabolic phenotype. This pattern suggests that severe metabolic dysregulation may better capture late-stage deterioration than inflammatory cytokine measures. An integrative schematic summarizing the constellation of clinical, inflammatory, and metabolic abnormalities observed in mouse *#15* is provided in Figure 2e.

## Discussion

This study illustrates that, even within a highly penetrant autoimmune model, an extreme systemic phenotype can emerge without a corresponding escalation in inflammatory activity. Although based on a single individual, documenting such an extreme presentation remains informative, highlighting biological divergence within a genetically homogeneous strain. These findings suggest that, at advanced disease stages, systemic inflammatory markers may reach a plateau beyond which they no longer discriminate severity, thereby limiting their capacity to capture divergent pathological trajectories [9, 10].

In contrast, metabolic profiling revealed a distinctly altered biochemical landscape that clearly set this mouse apart from all other. The magnitude and coherence of these metabolic abnormalities, including coordinated disruptions in urea-cycle–related nitrogen handling, sulfur amino-acid metabolism, and muscle-derived metabolites, indicate that downstream metabolic dysfunction may serve as a more discriminating correlate of extreme systemic deterioration. This aligns with observations in human lupus, where metabolic phenotypes often diverge independently of inflammatory indices [11–13], supporting the notion that metabolic dysregulation becomes a predominant dimension of disease heterogeneity associated with extreme disease severity once inflammation stabilizes at chronically elevated levels. In this context, metabolic readouts may offer greater sensitivity for detecting late-stage systemic deterioration than cytokine measurements alone.

Overall, this work underlines the importance of integrating metabolic assessments alongside inflammatory markers in autoimmune research. Extreme phenotypes can reveal mechanistic inflection points that remain concealed in group-averaged analyses, thereby enhancing the translational relevance of preclinical models. Although causality cannot be inferred from a single observation, the coherence of the clinical and metabolic alterations strengthens its value as a representative extreme phenotype. Such rare divergences highlight the importance of considering individual-level variation when interpreting autoimmune phenotypes.

## Author Contributions

KM: Conceptualization, data curation, formal analysis, investigation, visualization, writing – original draft, writing – review & editing. CJ: Investigation, writing – review & editing. DB: Investigation, writing – review & editing. AGMN: Investigation, writing – review & editing. HJD: Conceptualization, investigation, writing –review & editing.

